# Microevolution and phylogenomic characterization with perspectives in the 2022-2023 outbreak of child Respiratory Syncytial Virus type A

**DOI:** 10.1101/2023.01.28.526017

**Authors:** Sidra Majaz, Ashfaq Ahmad, Aamir Saeed, Shumaila Noureen, Faisal Nouroz, Amr Amin, Yingqiu Xie

**Author notes:** Equal contribution (Sidra Majaz, Ashfaq Ahmad, Aamir Saeed). Corresponding authors: Ashfaq Ahmad and Yingqiu Xie.

## Abstract

A communal respiratory syncytial virus (RSV) causes mild to severe illness, predominantly in older adults, or people with certain chronic medical conditions, and in particular, in young children. Symptoms may include runny nose, cough, fever, and difficulty breathing. In most cases, the infection is mild and resolves on its own, but in some cases, it can lead to more serious illness such as bronchiolitis or pneumonia. The RSV genome codes for ten proteins, NS1, NS2, N, P, M, SH, G, F, M2 and L. We aimed to identify the RSV geographical distribution and transmission pattern using site parsimonious frequencies, and investigate hotspot regions across the complete RSV genomes. These results indicated that RSV strains circulating in South and North America are not mixed to the European samples, however, genomes reported from Australia are the direct decedents of European samples. Samples reported from the United Kingdom were found diverse. Further, this report provides a comprehensive mutational analysis of all the individual RSV genes and in particular the 32 hotspot substituting regions circulating across the globe in RSV type A samples. This is the first comprehensive analysis of RSV type A that features mutational frequencies across the whole genome providing more clues for epidemiological control and drug development.

## Introduction

Viruses have become a new threat to mankind and thus drawn more attention due to their potential to cause large scale pandemic as we learned the lesson from COVID-19 [1]. Respiratory syncytial virus (RSV), also known as human respiratory syncytial virus (hRSV) contributes to the infection of respiratory tract [2]. Initially, RSV was isolated from Chimpanzee in 1956 and was also simultaneously recovered from infants with severe lower tract respiratory disease [3]. In clinical manifestations, RSV is linked to mild upper respiratory tract illness (URTI) or otitis media to severe and potentially life-threatening lower respiratory tract involvement (LRTI). The most prevalent form of LRTI in RSV-infected newborns is bronchiolitis, however there are reports indicating pneumonia and croup. Approximately, in infants and young children ~15–50 % of the lower airways were found affected in primary RSV infection that results in hospitalization and higher mortality [4, 5]. Apart from supportive care like fluid intake and rest, so far there is no specific treatment for RSV infection [6].

RSV is a non-segmented negative-sense single-stranded enveloped RNA virus that belongs to the family of Paramyxoviridae, genus Pneumovirus, and subfamily Pneumovirinae. The complete genome of RSV encodes ten proteins, M, M2, three envelope proteins, the fusion (F) glycoprotein, the G glycoprotein, and the small hydrophobic (SH) protein. Besides, five other structural and non-structural proteins are coded by the RSV genome, i.e., the large (L) protein, phosphoprotein (P), nucleocapsid (N), and non-structural proteins 1 and 2 (NS1 and NS2). Among them, protein G and F are important in host cell attachment, fusion and cellular entry [5, 7–9].

The RSV has been classified into two subtypes A and B which further includes strains like GA1 - GA7, SAA1, NA1 - NA4 and ON1 (RSV-A), and GB1 - GB4, SAB1 - SAB4, URU1 - URU2, BA1 - BA10, BA - C and THB (RSV-B) [10, 11]. There are few reports discussing mutations in individual genes like G, however we did not find any extensive analysis for RSV genomes. Here we are reporting for the first time all the mutational frequencies, hotspot regions and transmission of RSV subtype A. These analyses highlight a wider view of RSV transmission across different geological zones that could aid in predicting the oncoming pandemic and vaccine development. Besides transmission, this report provides all the observed mutations in genome coding regions and particularly the hotspot nucleotide sites prone to mutations in individual genes of RSV.

## Methodology

All the sequences used in these analyses were collected from the NCBI Virus database [12], where we selected taxonomic identification or taxid 208893, and spotted 11956 nucleotide sequences. Specific filters were applied for genome completeness, thus the final dataset we found contained 871 sequences. The whole dataset was exported to MAFFT for whole genome alignment [13]. Next, the aligned dataset was fed to the viral genome evolutionary analysis system (VENAS) for further analyses [14].

### Calculation for effective parsimony-informative site (ePIS) and network construction

To calculate ePIS, we followed a rule that the site will be considered parsimony-informative if it contains a minimum of two types of nucleotides in the aligned data, and at least two sites of them should occur with a minimum nucleotide frequency of two. Besides, the ePIS was considered effective only in the case that the site must contain unambiguous bases ≥80% of the total genomes. Keeping the above rules, 474 genomes were found satisfactory, and thus all the remaining analyses were carried out on the dataset containing 474 genomes. The derived ePIS results were used to classify all the genomes in haplotypes, and for this reason sequences containing similar ePIS were grouped onto the similar nodes and vice versa. Visualization, graph rendering and community detections were performed by Gephi.

### Mutational Analysis

Mutational analyses for individual genes were performed by NGS analysis package BioAider [12]. To retrieve codon information, aligned sets for individual genes were extracted from the aligned dataset used for VENAS. All the gaps (InDel operations) were removed and aligned them again using the codon alignment tool. The complete reports for individual genes substitutions can be found in table S1. To detect hotspot regions, only those non-synonymous substitutions were considered which showed changes in amino acid properties and more than 200 samples responded. The axillary art work was drawn by illustrator for the visualization of biological sequences (IBS) and Paint.net [13].

## Results

We retrieved the complete nucleotide sequence dataset for RSV type A (taxid. 1439707) from the NCBI Virus database. By December 2022, it contained 1166 nucleotide records, including 871 complete genomes. After initial filtration and name tagging that includes, genome ID, reported year and country, we applied viral genome evolutionary analysis system (VENAS) [14] and analyzed the evolutionary relationship between different genomes or RSV strains. Among them the earlier reported genome U39662.1 in 1997 was considered as a reference. Based on the effective parsimony informative sites (ePIS), and removal of redundant genome sequences, VENAS picked 474 genomes for further analyses. The final genome dataset contains sequences from the USA (2014, 2017, 2019, and 2021), Brazil (2021), Kenya (2021), Philippines (2017), Jordan (2018), Thailand (2019), Netherland (2021), Australia (2020, and 2022), China (2018), UK (2021), Spain (2021), Germany (2020), and four Unknown sequences (2022).

### RSV transmission distribution on the country scale

In order to trace the transmission routes, and contribution, we mapped 474 RSV genomes on the viral evolution network. Our data indicated individual clusters such as cluster of the USA 2014 sequences, connected to cluster of the USA 2017, which further connected the Brazilian cluster 2021, whereas sequences from Jordan 2018 and Philippines 2017 were also observed. Apparently, we found that samples of North and South America were cluster separated from the European samples. However, genomes reported from the USA 2019 were found mixed with genomes reported from Europe (Figure 1A).

**Figure 1.**
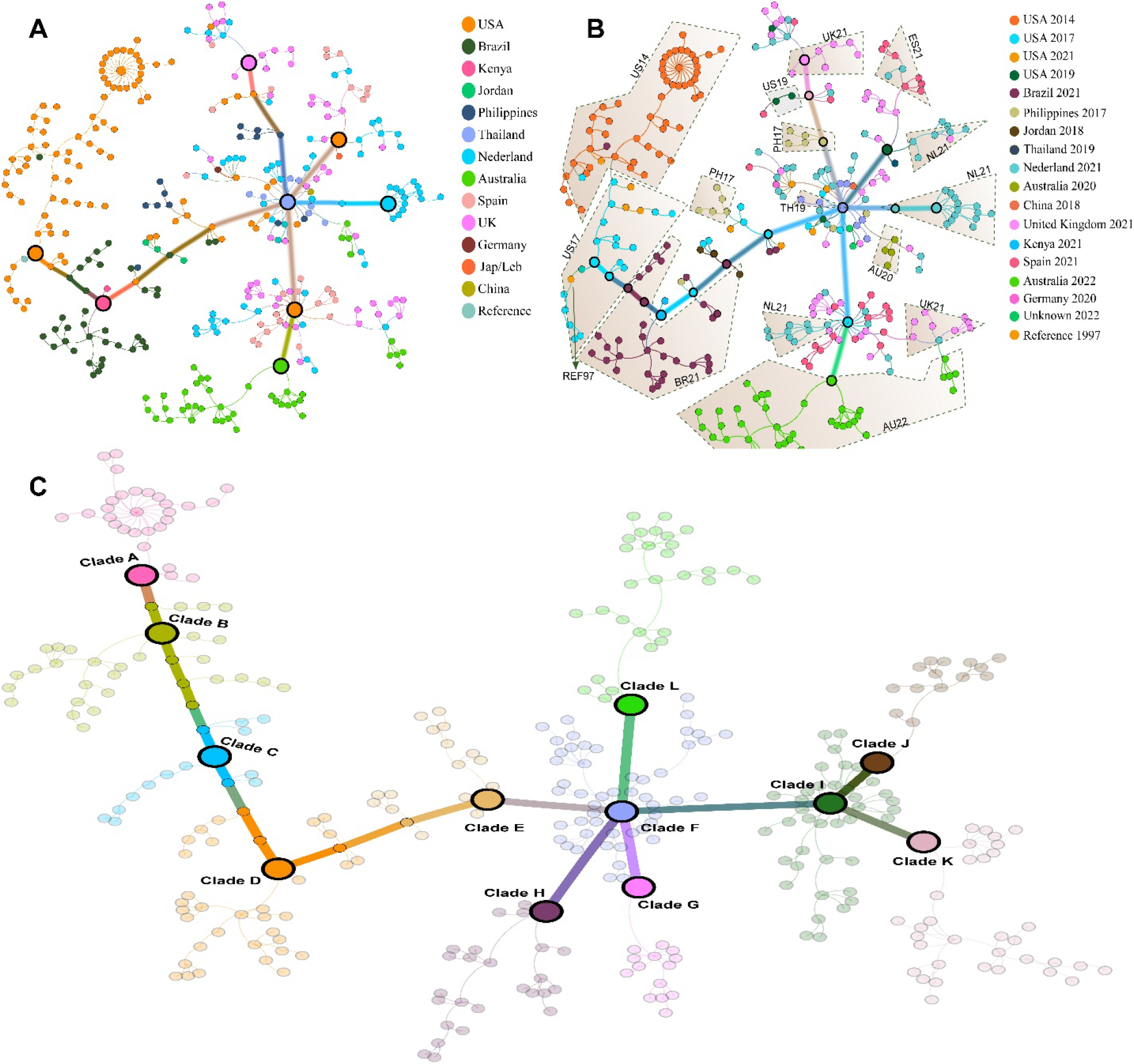
RSV genome distribution. (A), Country wise network distribution of the RSV genomes derived through effective parsimony informative sites (ePIS). (B), Country and year wise network of complete RSV genomes. (C), Calculated transmission pattern via community detection by modularity approach.

To understand the transmission pattern, we used a network modularity function and clustered the whole network beyond their country or year-wise taglines. We retrieved 12 clades (clade A to clade L). Interestingly, our data predicted a unique pattern, for instance the terminal clade A, contained purely genomes reported from the USA in the year 2014, which connected clade B retaining genomes reported from the USA in the year 2014 and 2017. Further, clade C, a direct descendent of clade B contains all the genomes reported in 2017, except one sequence from the USA 2021. Apart from few genomes reported from the USA 2017, Philippines 2017 and Jordan 2018, clade D contain > 95% of the genomes reported from Brazil 2021. To our interest the smaller clade E, contains representation of almost all the previous clades and connects the bigger node clade F. All the previous clades did not show a single genome reported from the Europe therefore, for better understanding, we call clade F, a European clade, as it is the first with European representation that also gave birth to diverse nodes reported from European countries.

The European node (clade F), is tetra-furcated into clade G, H, I and L. Among them, clade H contained samples of Nederland, Spain and the United Kingdom reported in 2021 while clade G was found enriched with all the sequences reported from Nederland 2021. Likewise, clade L is formed from the genomes reported in the United Kingdom 2021, the USA 2019, and two genomes from Spain 2021. Interestingly, clade L emerged from the European cluster through genomes reported from Philippines 2017 and the USA 2019. Finally, Clade I that contains genomes from Spain, Nederland and the United Kingdom 2021 which is bifurcated to clade J and K, where clade J contains samples from the United Kingdom 2021 that leads to Australia 2022 and the clade K only contains all the genomes reported from Australia reported in 2022. All the samples are mapped with country, year and transmission wise in (Figure 1BC).

Collectively, these analyses indicate the global prevalence and the presence of different RSV type A strains. For instance, genomes reported from Australia in 2022 were gathered in two different clusters emerging from the European genomes, suggesting the presence of two different strains. However, genomes reported from Australia in 2020, are lying distantly for those reported in 2022. Likewise, genomes from the UK reported in 2021, were found almost everywhere with European genomes, depicting the possibility of numerous RSV strains. Genomes from Spain and the Netherlands were also found in 2 and 3 different nodes, highlighting the circulation of more than one strain in that country.

### Mutation and substitution frequencies of synonymous and non-synonymous sites in RSV genomes

According to the NCBI records, RSV contains ten to eleven protein coding genes, i.e., non-structural protein 1 and 2 (NS1 and NS2), nucleoprotein (N), phosphoprotein (P), matrix 1 protein (M), small hydrophobic protein (SH), attachment protein (G), fusion protein (F), matrix 2 protein (M2), and polymerase (L). We calculated substitution observed in RSV genomes for all coding sequences (CDS) of all ten genes, particularly synonymous and non-synonymous substitutions. All the substitutions were accessed and calculated against the reference strain U39662.1. A total of 6257 (45%) sites were observed in substitutions, and among them 2099 (15.1%), 3027 (21.7%), 611 (4.3%), and 473 (3.4%) were synonymous, non-synonymous, both and terminations, respectively. Among the non-synonymous substitutions, 1442 (10%) sites were found to have changes in amino acid properties. The highest termination substitutions were found in L followed by N and G proteins (Table. 1).

**Table 1.**
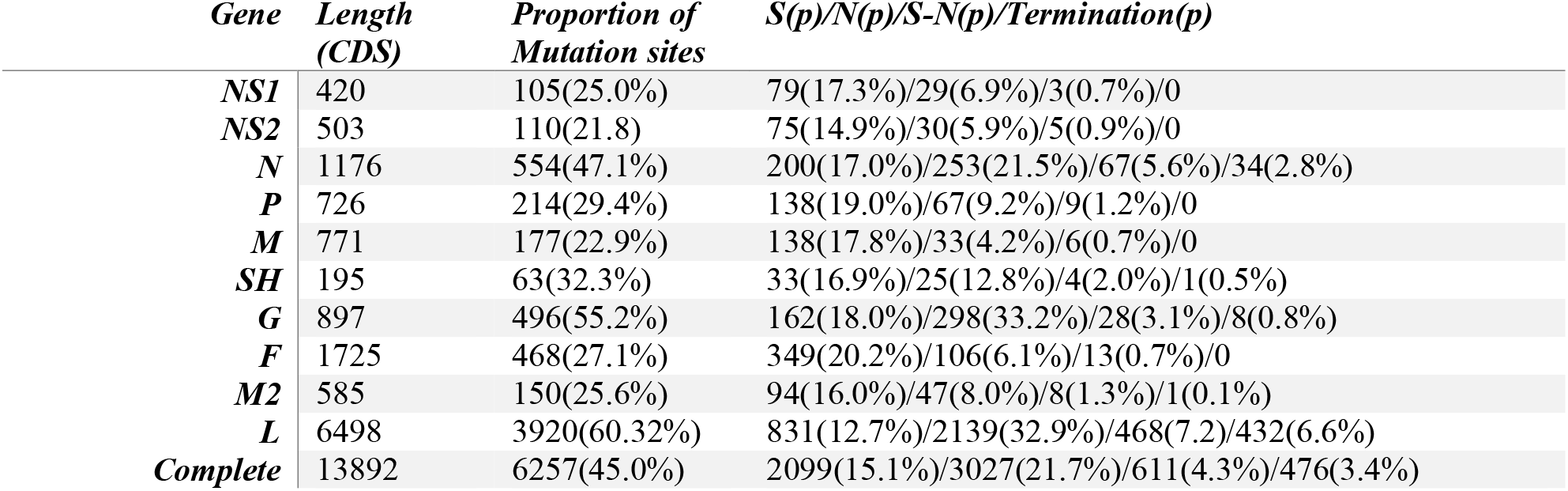
Statistics of all types substitutions observed in individual genes of RSV

| *Gene* | *Length*<br>*(CDS)* | *Proportion of*<br>*Mutation sites* | *S(p)/N(p)/S-N(p)/Termination(p)* |
| --- | --- | --- | --- |
| *NS1* | 420 | 105(25.0%) | 79(17.3%)/29(6.9%)/3(0.7%)/0 |
| *NS2* | 503 | 110(21.8) | 75(14.9%)/30(5.9%)/5(0.9%)/0 |
| *N* | 1176 | 554(47.1%) | 200(17.0%)/253(21.5%)/67(5.6%)/34(2.8%) |
| *P* | 726 | 214(29.4%) | 138(19.0%)/67(9.2%)/9(1.2%)/0 |
| *M* | 771 | 177(22.9%) | 138(17.8%)/33(4.2%)/6(0.7%)/0 |
| *SH* | 195 | 63(32.3%) | 33(16.9%)/25(12.8%)/4(2.0%)/1(0.5%) |
| *G* | 897 | 496(55.2%) | 162(18.0%)/298(33.2%)/28(3.1%)/8(0.8%) |
| *F* | 1725 | 468(27.1%) | 349(20.2%)/106(6.1%)/13(0.7%)/0 |
| *M2* | 585 | 150(25.6%) | 94(16.0%)/47(8.0%)/8(1.3%)/1(0.1%) |
| *L* | 6498 | 3920(60.32%) | 831(12.7%)/2139(32.9%)/468(7.2)/432(6.6%) |
| *Complete* | 13892 | 6257(45.0%) | 2099(15.1%)/3027(21.7%)/611(4.3%)/476(3.4%) |

In protein L and G, we also observed higher and similar non-synonymous mutation frequency compared to all other genes of RSV. However, F and P proteins have relatively shown a higher number of synonymous substitutions than non-synonymous. Complete details for substitutions and substitution type for all individual genes can be accessed in form (Table S1). To evaluate the overall substitution frequency of the mutated sites, present in all ten CDSs, we alienated the substitution frequency into seven different groups (G1 to G7), and depicted the frequency distribution of 5126 substituted sites (3027 non-synonymous and 2097 synonymous) of the 474 sequenced genomes. Group number on the X-axis indicates the number of strains participating in a substitution event at a particular site, whereas the Y-axis shows the number of substitutions in a respective CDS (Figure 2).

**Figure 2.**
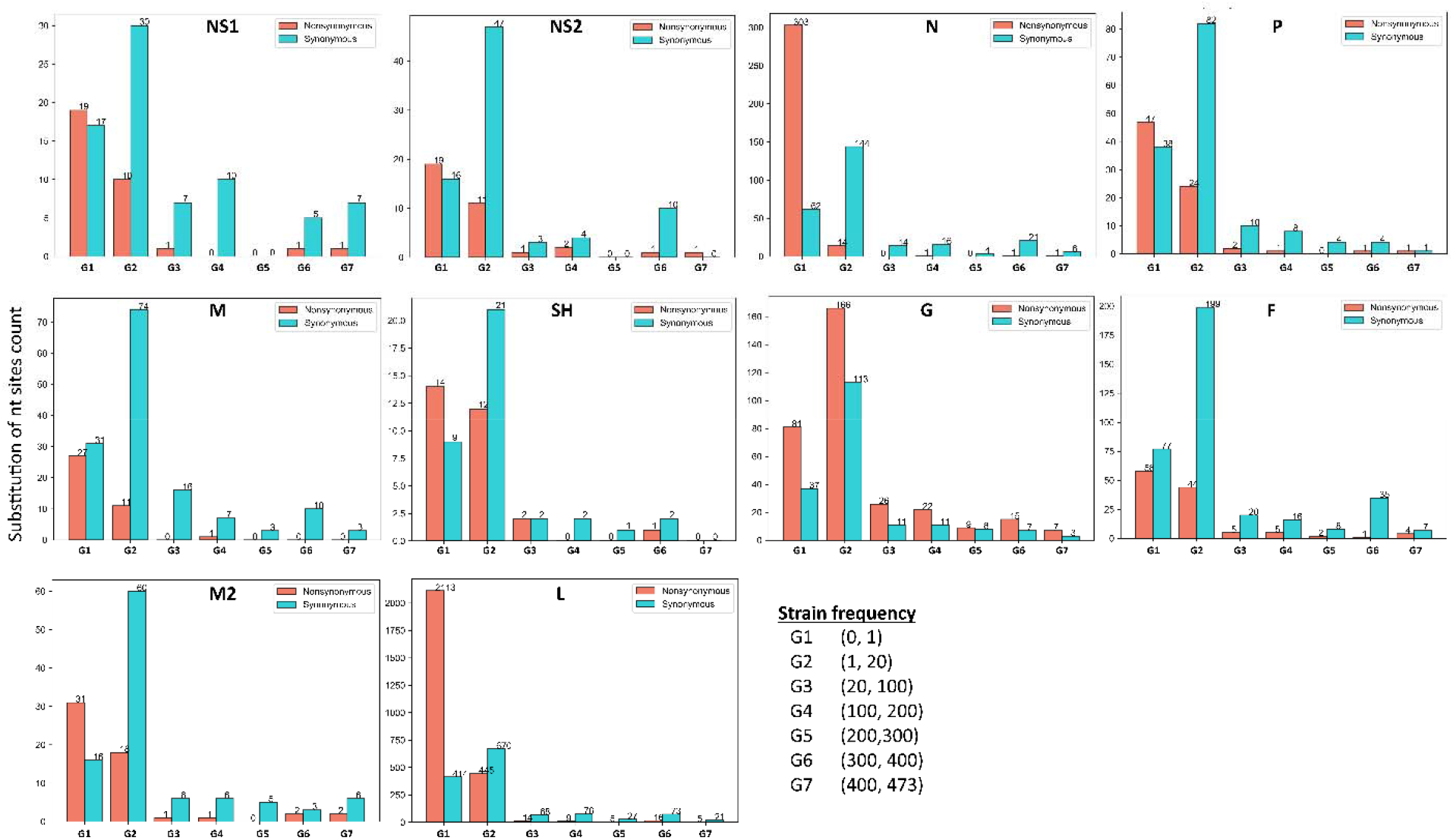
Overall substitution frequencies of all coding genes coded by the RSV genome. Frequencies of only synonymous and non-synonymous substitutions are shown here. Y-axis represent the number of substitutions and the X-axis depicts number of genomes or samples participated.

Our results indicated that apart from the initial groups, where non-synonymous mutations were found relatively higher than the distribution of synonymous mutations. Comparing all the CDSs, the CDS of G protein showed the highest number of non-synonymous mutations where 55 mutations were found in G4 - G7, meaning the participation of 200 samples. We have also observed that CDSs of NS1, NS2, SH, and M proteins offered relatively lower sites for non-synonymous mutations in the majority of the sequenced RSV strains. The distribution of substitution frequency of each codon in each gene can be found in the supplementary Table. Overall, these analyses provide complete mutational information of all ten RSV’s CDSs. Such key features can also be used to assess the evolutionary pressure on selected sites or response to therapeutic agents.

### Substitution hot-spots in RSV type A genomes

Next, we were interested to identify the hot-spot substitution sites in RSV CDSs. Similar to the analysis [15] for SARS-CoV-2, we defined a criterion for hotspot regions. A site with a substitution frequency over 200 will be considered a hot-spot site, or a site that offered substitution to more than 200 RSV strains (42% in our case) will be considered a potential substitution hot-spot. Second, the respective site must allow non-synonymous substitution and the observed amino acid should record a change in amino acid properties. A total of 367 substitution sites were found with > 200 substitution frequency where 290, and 77 were synonymous and non-synonymous respectively. Among 77 non-synonymous substitution sites, 32 were those affecting amino acid properties (either changing polarity or charge difference) were considered hot-spot substitution sites. These 32 hot-spots were found distributed across F protein (4), G protein (13), L protein (10), N1, N2, P, N, and M2 (1 each) (Figure 3). Two hot-spots that resulted in CAA to TTA (Q142L) in G protein and ATA to ACG and ACA (I184T) in M2 protein offered double substitutions.

**Figure 3.**
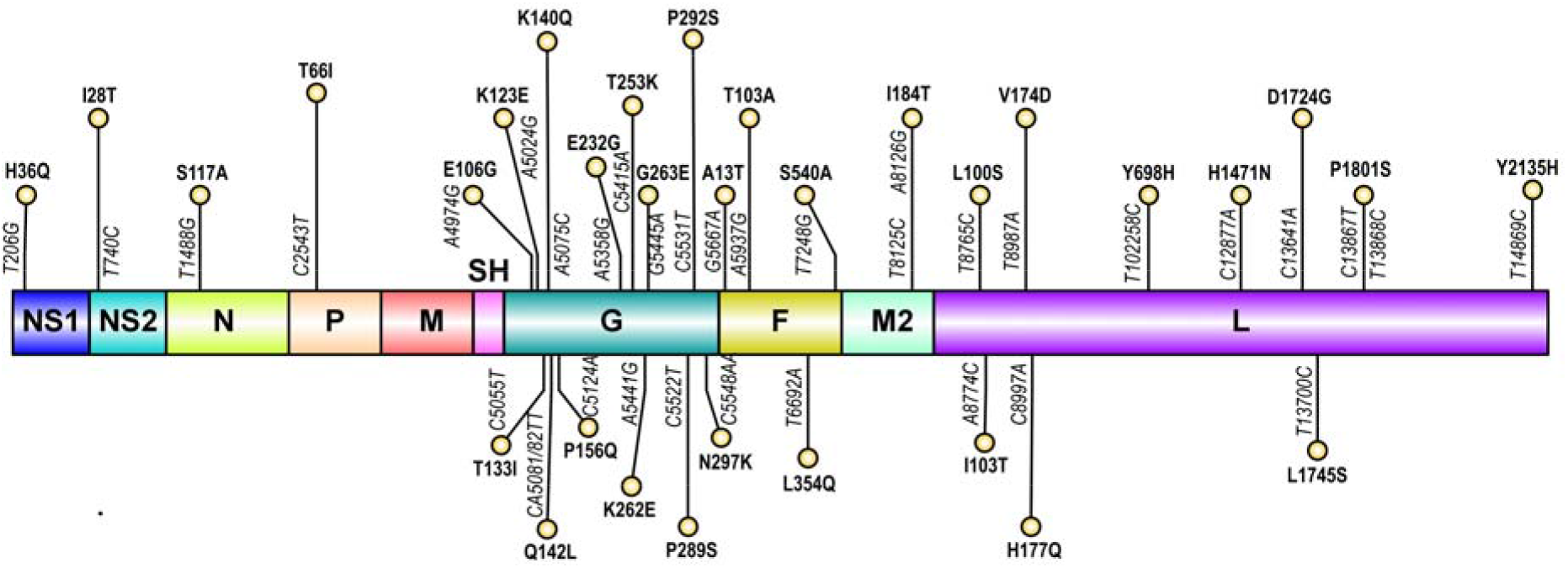
Presentation of hotspots mutations in all proteins. The nucleotide positions are numbered according to the reference genome U39662.1 while the proteins are numbered as per protein.

### Discussion

Respiratory syncytial virus (RSV) is a communal respiratory virus that can cause mild to severe illness, predominantly in young children, elderly adults, and people with certain chronic medical conditions. In some cases, it can lead to more serious illness such as bronchiolitis or pneumonia. Here, we detected a probable transmission pattern that in turn may also explain the classification of different strains circulating worldwide. Reference to the hotspot substitutions, we identified 32 hotspots positions, where 13 and 4 are detected only the G and F protein. Both G and F proteins are considered important because both can induce neutralizing antibodies, and are heavily glycosylated, which has been shown to affect with antibody recognition [17-19]. Structurally, the G protein comprises three domains; a cytoplasmic domain (1–37 amino acids), a transmembrane domain (38-66 amino acids), and an ectodomain region (67–312) [20, 21]. Interestingly, all the hotspot positions we identified belong to the ectodomain of the G protein. There are individual reports in G, F, and L proteins [22-27] and we believe that this report will assist researchers to directly pick the hotspot regions for biochemical testing.

### Conclusion

This report provides an idea to probe the presence of any viral species and classify them into different strains. Such approaches can be further utilized towards personalized medication therapy particularly in children to cope pandemic.

## Authors Contribution

i. Experiment performed and data analysis by SM, AA, AS
ii. Idea conceived and refined by AA, YX
iii. Manuscript write up were performed by SM, AA, AS and AAm
iv. Art work was generated by AA, AS
v. Critical reading and directions were given by AA, AAm, YX
vi. Funding support and/or supervision by AA, AAm, YX

